# *Drosophila* models of PIGA-CDG mirror patient phenotypes

**DOI:** 10.1101/2023.10.27.564441

**Authors:** Holly J. Thorpe, Katie G. Owings, Miriam C. Aziz, Madelyn Haller, Emily Coelho, Clement Y. Chow

## Abstract

Mutations in the phosphatidylinositol glycan biosynthesis class A (PIGA) gene cause a rare, X-linked recessive congenital disorder of glycosylation (CDG). PIGA-CDG is characterized by seizures, intellectual and developmental delay, and congenital malformations. The *PIGA* gene encodes an enzyme involved in the first step of GPI anchor biosynthesis. There are over 100 GPI anchored proteins that attach to the cell surface and are involved in cell signaling, immunity, and adhesion. Little is known about the pathophysiology of PIGA-CDG. Here we describe the first *Drosophila* model of PIGA-CDG and demonstrate that loss of *PIG-A* function in *Drosophila* accurately models the human disease. As expected, complete loss of *PIG-A* function is larval lethal. Heterozygous null animals appear healthy, but when challenged, have a seizure phenotype similar to what is observed in patients. To identify the cell-type specific contributions to disease, we generated neuron- and glia-specific knockdown of *PIG-A*. Neuron-specific knockdown resulted in reduced lifespan and a number of neurological phenotypes, but no seizure phenotype. Glia-knockdown also reduced lifespan and, notably, resulted in a very strong seizure phenotype. RNAseq analyses demonstrated that there are fundamentally different molecular processes that are disrupted when *PIG-A* function is eliminated in different cell types. In particular, loss of *PIG-A* in neurons resulted in upregulation of glycolysis, but loss of *PIG-A* in glia resulted in upregulation of protein translation machinery. Here we demonstrate that *Drosophila* is a good model of PIGA-CDG and provide new data resources for future study of PIGA-CDG and other GPI anchor disorders.

**Article Summary:** PIGA-CDG is a rare genetic disorder. In order to study this rare disease, we generated and characterized several Drosophila models of PIGA-CDG. These models faithfully recapitulate different patient phenotypes, including movement disorder and seizures. Drosophila is a good model for PIGA-CDG and other GPI anchor disorders.

## Introduction

PIGA-CDG (phosphatidylinositol glycan biosynthesis class A Congenital Disorder of Glycosylation) is an ultra-rare, X-linked recessive disorder caused by partial loss of function mutations in the *PIGA* gene (JOHNSTON *et al*. 2012; KATO *et al*. 2014; VAN DER CRABBEN *et al*. 2014; BAYAT *et al*. 2020). Patients with PIGA-CDG display a range of symptoms affecting many systems including neurological abnormalities, muscular abnormalities, and skeletal abnormalities, with most patients presenting with seizures, hypotonia, and neurodevelopmental delay. There are fewer than 100 reported patients with PIGA-CDG with ∼40 different mutations (BAYAT *et al*. 2020). The underlying mechanisms of PIGA-CDG are not understood, and current treatment options are limited. Most treatments for CDGs focus on alleviating symptoms, rather than correcting the cause of the disorder.

Glycosylphosphatidylinositol (GPI) anchor synthesis is a highly conserved pathway involving over 30 proteins which builds a sugar chain on a phosphatidylinositol molecule in a stepwise manor. The *PIGA* gene encodes phosphatidylinositol glycan biosynthesis class A, the catalytic enzyme involved in the first step GPI anchor synthesis (MIYATA *et al*. 1993; WATANABE *et al*. 1998; KINOSHITA AND INOUE 2000). The first steps of GPI anchor biosynthesis occur on the cytoplasmic side of the endoplasmic reticulum (ER). In the first step, PIGA transfers an N-acetylglucosamine (GlcNAc) from uridine 5′-diphospho N-acetylglucosamine (UDP-GlcNAc) to an existing phosphatidylinositol (PI) within the ER membrane, generating the first intermediate of GPI anchor synthesis, N-acetylglucosaminyl phosphatidylinositol (GlcNAc-PI).

GPI anchors attach over 100 proteins to the surface of the cell that are involved in many functions including cell signaling, immunity, and adhesion. Loss of PIGA function leads to decreased surface expression of GPI anchored proteins (LUKACS *et al*. 2020; LIU *et al*. 2021). Typical PIGA-CDG patients display 5-15% of normal GPI anchored proteins on the cell surface (BAYAT *et al*. 2020). GPI anchored protein precursors have a C-terminal signal sequence that is normally cleaved when the protein is attached to the anchor (KINOSHITA 2020). If GPI anchor synthesis is inhibited before addition of the first mannose to the anchor (such as in PIGA-CDG), the C-terminal sequence is not cleaved from the protein, and the protein is recognized as a misfolded protein and sent to be degraded by the proteasome (KINOSHITA 2020). It is unclear if the symptoms observed in PIGA-CDG are due to cellular stress caused by the misfolded proteins or if they are due to loss of specific GPI anchored proteins, although it is likely due to a combination of both factors.

Even though the first reported PIGA-CDG patient was published in 2012 (JOHNSTON *et al*. 2012), there have only been a handful of cellular or animal model studies. Cells with *PIGA* null mutations are viable, but show decreased abundance of GPI anchored proteins on the surface of the cell (LUKACS *et al*. 2020; LIU *et al*. 2021). While cell culture systems are useful for investigating the cellular effects of loss of *PIGA*, many effects are missed in cell models because there are many cell-type specific GPI anchored proteins. Mouse models have also been used to investigate PIGA-CDG (KAWAGOE *et al*. 1996; LUKACS *et al*. 2019; LUKACS *et al*. 2020; KANDASAMY *et al*. 2021; JANGID *et al*. 2022). Global knockout of *PIGA* in mice is embryonic lethal (KAWAGOE *et al*. 1996). Conditional knockout of *PIGA* in the mouse central nervous system results in ataxia and degenerative tremors (LUKACS *et al*. 2020). To date, no *Drosophila* models have been reported.

Here, we generated *Drosophila* models to investigate the pathophysiology of phenotypes observed in PIGA-CDG. Homozygous *PIG-A* null alleles result in larval lethality, but the heterozygotes are viable and have a seizure phenotype similar to what is observed in patients. Cell specific knockdowns (KD) in glia and neurons lead to distinct neurological phenotypes. KD in the neurons causes neuromuscular defects while KD in glia results in seizures. Transcriptome analyses on these different models showed distinct transcriptomic signatures when PIGA function is reduced in different cell types. Together, this is the first report of a *Drosophila* model of PIGA-CDG and will be a rich resource for future studies on the pathogenesis of this understudied rare disorder.

## Materials and Methods

### *Drosophila melanogaster* fly stocks

All stocks were maintained under standard laboratory conditions on agar–dextrose–yeast medium at 24°C on a 12-h light/dark cycle. The following strains are from Bloomington *Drosophila* Stock Center: *PIG-A* RNAi (62696), repo-GAL4 (7415), and elav-GAL4 (46655). The corresponding attP40 control strain was used as our wild-type control. The *PIG-A* null allele (*PIG-A*^*-/-*^ and *PIG-A*^*+/*-^) was generated as a *PIG-A* Kozak-miniGAL4 allele (KANCA *et al*. 2019) using the sgRNAs TATGTGGTATGTCAAATTACTGG and TTGGCTAGTTGATGGAAGAATGG.

### qPCR

Total RNA was extracted from tubulin-GAL4 > PIG-A RNAi (Bloomington *Drosophila* Stock Center: 62696) and control larvae using TRIzol Reagent (ThermoFisher Cat #15596026) followed by Direct-zol RNA Miniprep with DNAse step (Zymo Research R2051). Each biological replicate contained 15-20 larvae. 1ug of RNA was used to synthesize cDNA using the ProtoScript® II First Strand cDNA Synthesis Kit (NEB Cat #E6560L). 50ng of cDNA was used to perform qPCR with PowerUp SYBR Green Master Mix (ThermoFisher Cat # A25741). All protocols were followed according to the manufacturer’s instructions. qPCR was run on a QuantStudio 3. Fold change of gene expression was calculated using the Delta-Delta Ct method. Primers used are from FlyPrimerBank (Primer ID #PP29152).

### Phenotypic analyses

#### Lifespan

Lifespan was monitored daily. 10 flies were placed in each vial. All flies were kept at 25 deg and maintained on a 12:12 light dark cycle. Flies were monitored for death once a day until all flies were dead. Survival analysis was performed using the Survival package in R (R version 4.2.0; survival package version 3.3-1; running under Windows 10 x64).

#### Climbing

Flies were maintained under standard conditions for at least 1 day after CO2 collection. All flies that were tested were 3-5 days of age. Ten flies were placed in empty vials and allowed to rest for 10 min. Vials were tapped to drop all flies to the bottom and climbing rate was measured.

#### Bang sensitivity

Flies were maintained under standard conditions for at least 1 day after CO2 collection. All flies that were tested were 3-5 days of age. Ten flies were placed in empty vials and allowed to rest for 10 min. Vials were vortexed at full speed for 10 seconds and seizure recovery was observed. Flies were considered to be seizing until they are upright and walking around.

### RNAseq

mRNA sequencing was performed on total RNA from heads of male control or knockdown 3-day-old flies. Neuron- and glial-specific knockdown had their own controls. All groups were sequenced in triplicate, for a total of 12 samples.

RNA was extracted using a Direct-zol RNA Miniprep (Zymo Research R2061) using TRIzol Reagent (ThermoFisher Cat # 15596026) and including the DNAse step. Samples were prepared and sequenced by the Huntsman Cancer Institute High-Throughput Genomics Core. Samples were sequenced on the NovaSeq 50 × 50 bp Sequencing, for ∼25 million paired reads per sample. Fastq files were trimmed using seqtk v1.2 software (for Fastq and processed files see GEO repository: GSE241512). RNAseq reads were aligned to the *Drosophila melanogaster* reference genome (assembly BDGP Release 6) using Bowtie2 v2.2.9 software (LANGMEAD AND SALZBERG 2012)and alignment files were sorted and converted using Samtools v1.12 (LI *et al*. 2009). Read counts were normalized using the default normalization method in DESeq2 (LOVE *et al*. 2014) package in R. Differential gene expression was assessed using linear models in the DESeq2 package. Genes were considered significantly differentially expressed if the adjusted p-value ≤ 0.05 and the fold change magnitude was ≥ 1.5 (Log2 fold change ≥ 0.585 or ≤ -0.585). GO analyses was performed using standard tools at www.geneontology.org.

## RESULTS

### Complete loss of *PIG-A* is lethal

We obtained a null allele of *PIG-A*, wherein the coding sequence is replaced by a GAL4 cassette (KANCA *et al*. 2019). Homozygous *PIG-A*^*-/-*^ adult flies are never observed. To determine when the *PIG-A*^*-/-*^ larvae die, we tracked their growth during larval development (Figure 1A). *PIG-A*^*-/-*^ are indistinguishable from controls in the 3^rd^ instar larval stage, up to 96 hrs. after egg laying. Beginning around this time, *PIG-A*^*-/-*^ larval growth slows and is significantly smaller than controls. *PIG-A*^*-/-*^ larvae never progress to pupation and languish as 3^rd^ instar, while wildtype flies pupate and eclose as adults.

**Figure 1.**
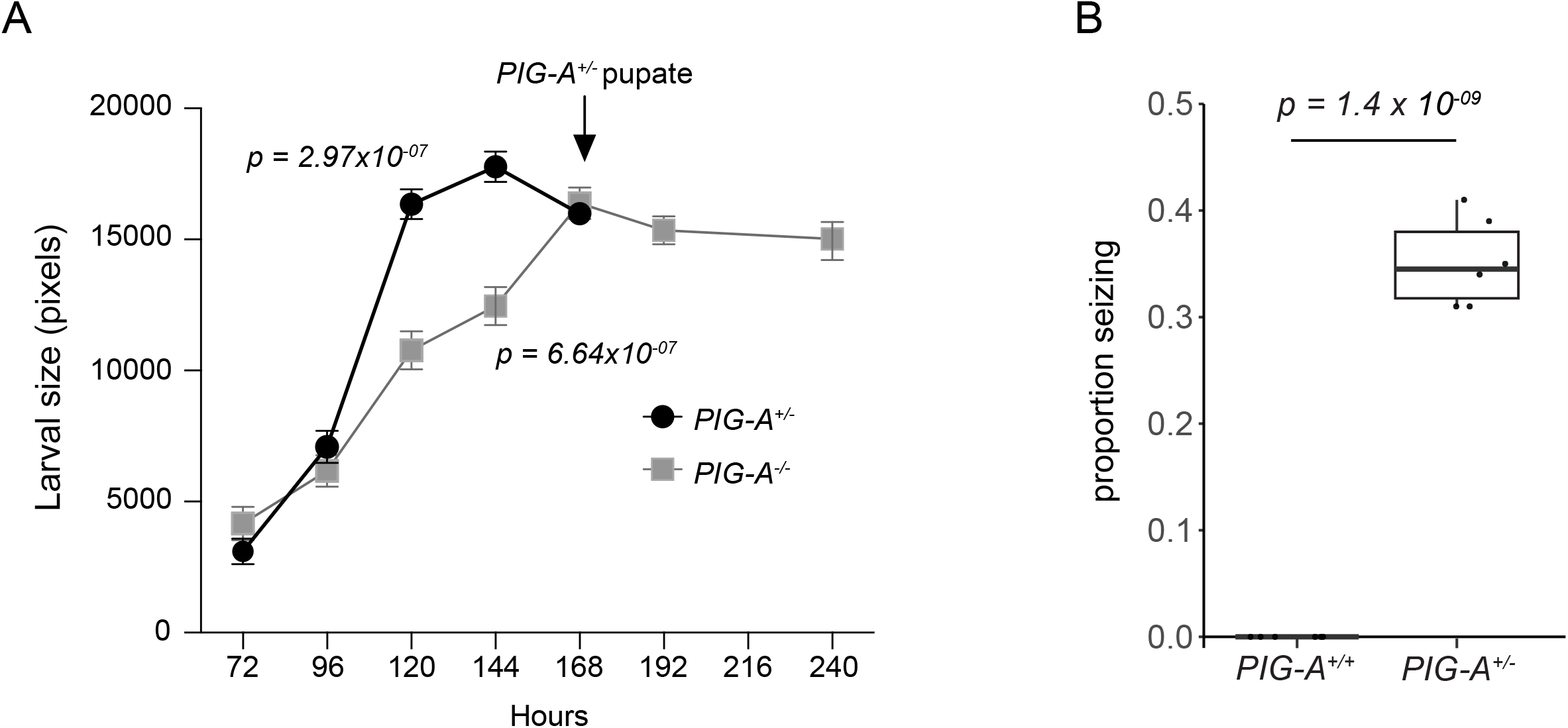
*PIG-A* null phenotypes. A) *PIG-A*^*-/-*^ larvae are smaller than *PIG-A*^*+/-*^ larvae and they fail to eclose. B) ∼35% of *PIG-A*^*+/-*^ flies display seizures upon bang sensitive testing. Each point represents a single experiment with at least 30 flies. Wildtype flies do not have seizures.

We also evaluated ubiquitous knockdown of *PIG-A* using a tubulin-GAL4 driver to express a UAS-RNAi (Figure 1B). We also never observed adult knockdown flies. Unlike the null flies, the ubiquitous KD pupae are indistinguishable from wildtype controls and do not die during development. Despite this, the ubiquitous KD animals still all die during pupation. The slight difference in survival and development suggests that the RNAi knockdown is not a complete knockdown, and some residual expression remains. In line with this, qRT-PCR of *PIG-A* transcript in ubiquitous KD larvae indicate that there is significant reduction in expression, with ∼39% residual transcript (39.6 ± 5.9; p < 0.0001) (Figure S1).

### Haploinsufficiency of *PIG-A* results in seizures

*PIG-A*^*+/-*^ flies develop normally and eclose as adults at the expected Mendelian ratio. However, we reasoned that these heterozygous null flies are more similar to PIGA-CDG patients than the homozygous nulls. Because PIGA is X-linked in humans, all reported PIGA-CDG patients carry a single partial loss-of-function mutations resulting in 50% or less activity (JOHNSTON *et al*. 2012; BAYAT *et al*. 2020). To evaluate if the *PIG-A*^*+/-*^ flies could model patient phenotypes and have neurological deficits, we first tested them for climbing ability, a general measure of nervous system function. The standard protocol for evaluating climbing ability entails tapping the flies to the bottom of the vial and measuring how long it takes to climb to the top. Upon tapping down these flies, we noticed that a subset would display seizure like behavior, preventing any meaningful climbing assessment. To formally test if *PIG-A*^*+/-*^ flies have seizures, we performed the bang sensitive assay, a commonly used measure of seizure susceptibility in flies. We found that on average, 35% of *PIG-A*^*+/-*^ flies have seizure activity following bang sensitive testing, compared to zero seizures in the WT flies (0.35 ± 0.04; p = 1.4 × 10^−9^) (Figure 2). This suggests that the heterozygous null flies may model some of the prominent patient phenotypes.

**Figure 2.**
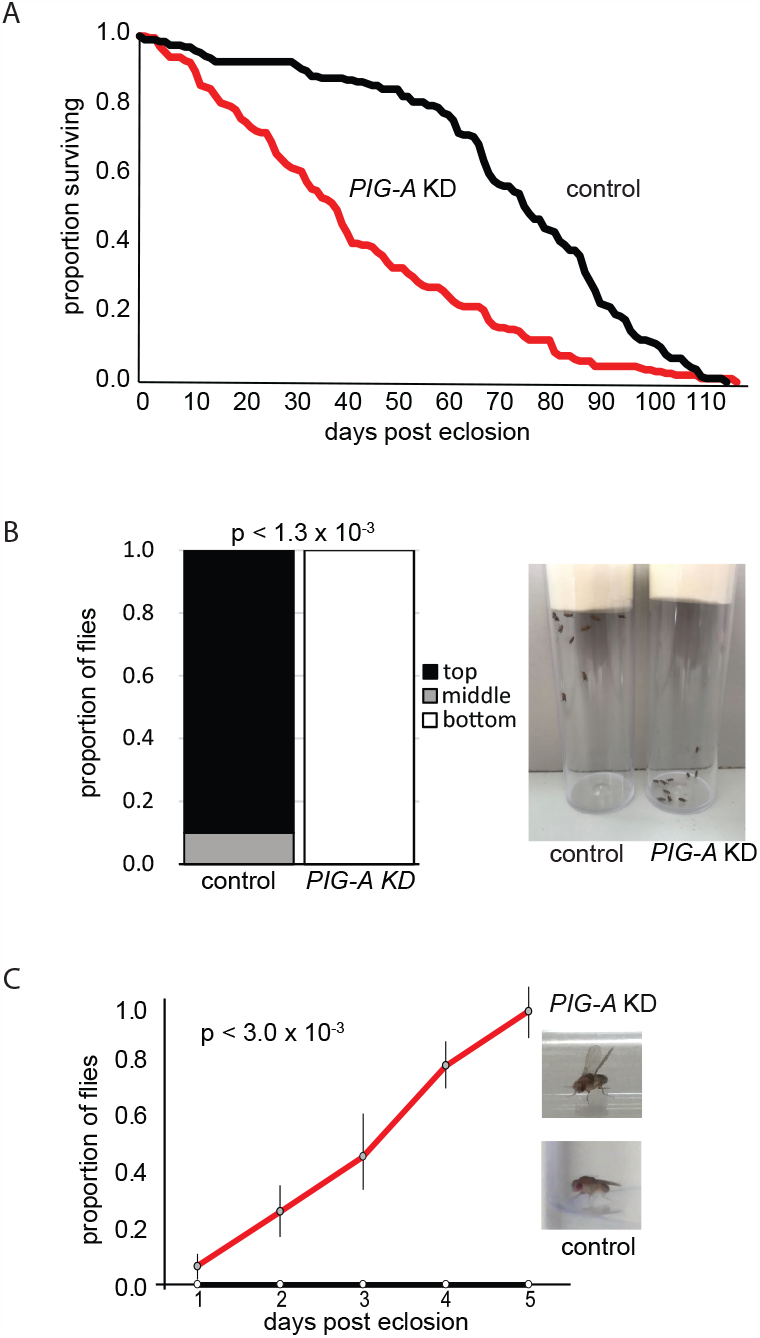
Phenotypes associated with neuron-specific knockdown of *PIG-A*. Flies with neuron-specific knockdown of *PIG-A* A) have reduced lifespan compared to control wildtype flies, B) fail to climb, and C) develop a degenerative erect wing phenotype. control = wildtype; *PIG-A* KD = neuron-specific knockdown

### Cell-type specific knockdown of *PIG-A*

Because most of the prominent phenotypes associated with PIGA-CDG are neurological (BAYAT *et al*. 2020), and to begin to uncover the cellular origins of the phenotypes, we performed neuron- or glial-specific knockdown of *PIG-A* to model neurological phenotypes observed in patients. To generate neuron-specific knockdown of PIGA, we used the pan-neuronal driver, elav-GAL4, to drive *PIG-A* RNAi (same as described above) in all neurons. Neuron-specific knockdown flies eclosed at expected Mendelian ratio but have a shorter lifespan than control flies (p < 2.0 × 10^−16^) (Figure 2A). Fifty percent of neuron-specific knockdown flies are dead by 40 days post eclosion, while more than 90% of control flies are still living. Neuron-specific knockdown flies display a climbing defect where nearly 100% of flies fail to climb to the top of vial in 20 seconds (Figure 2B). Control flies easily climb to the top in the same time interval (p < 1.3 × 10^−3^). Neuron-specific knockdown flies also display a degenerative erect wing phenotype (Figure 2C). They eclose with normal wing posture, but over five days, all the flies develop an erect wing phenotype. Control flies do not display a wing phenotype over this same period (p < 3.0 × 10^−3^). Strikingly, neuron-specific knockdown flies do not show a seizure phenotype upon treatment with a bang sensitive protocol.

To generate glial-specific knockdown of *PIG-A*, we used the pan-glial driver, repo-GAL4 to drive *PIG-A* RNAi in all glia. Glia-specific knockdown flies also eclosed at an expected Mendelian ratio but have a shorter life span than control flies. Fifty percent of glia-specific knockdown flies are dead at 60 days post eclosion, while nearly 80% of control flies are living (p < 2.0 × 10^−16^) (Figure 3A). Glia-specific knockdown flies have a slower rate of death than neuron-specific knockdown flies. Glia-specific knockdown flies have a severe seizure phenotype upon bang sensitivity testing (Figure 3B). Seventy percent of glia-specific knockdown flies display a very severe seizure phenotype where the flies completely pass out for nearly a minute before recovery. Fewer than 1% of control flies show a seizure phenotype with the same testing. This is likely an underestimate of the proportion of flies that seize because it appears that seizures can easily be elicited in glia-specific knockdown flies. We often observe flies that have passed out in a vial that has not been subject to bang sensitive testing. We also observe flies that pass out when someone walks by the vials. Thus, the 70% seizure rate is an underestimation because at any time, some of the flies are likely in a refractory period after a non-elicited seizure. Because of this severe seizure phenotype, we were unable to assay movement related deficits. Finally, ∼60% of glia-specific knockdown flies have a wrinkled, partially inflated wing phenotype, compared to normal wings in 100% of control flies (Figure 3C-D). The remaining ∼40% of glia-specific knockdown flies have wildtype normal appearing wings.

**Figure 3.**
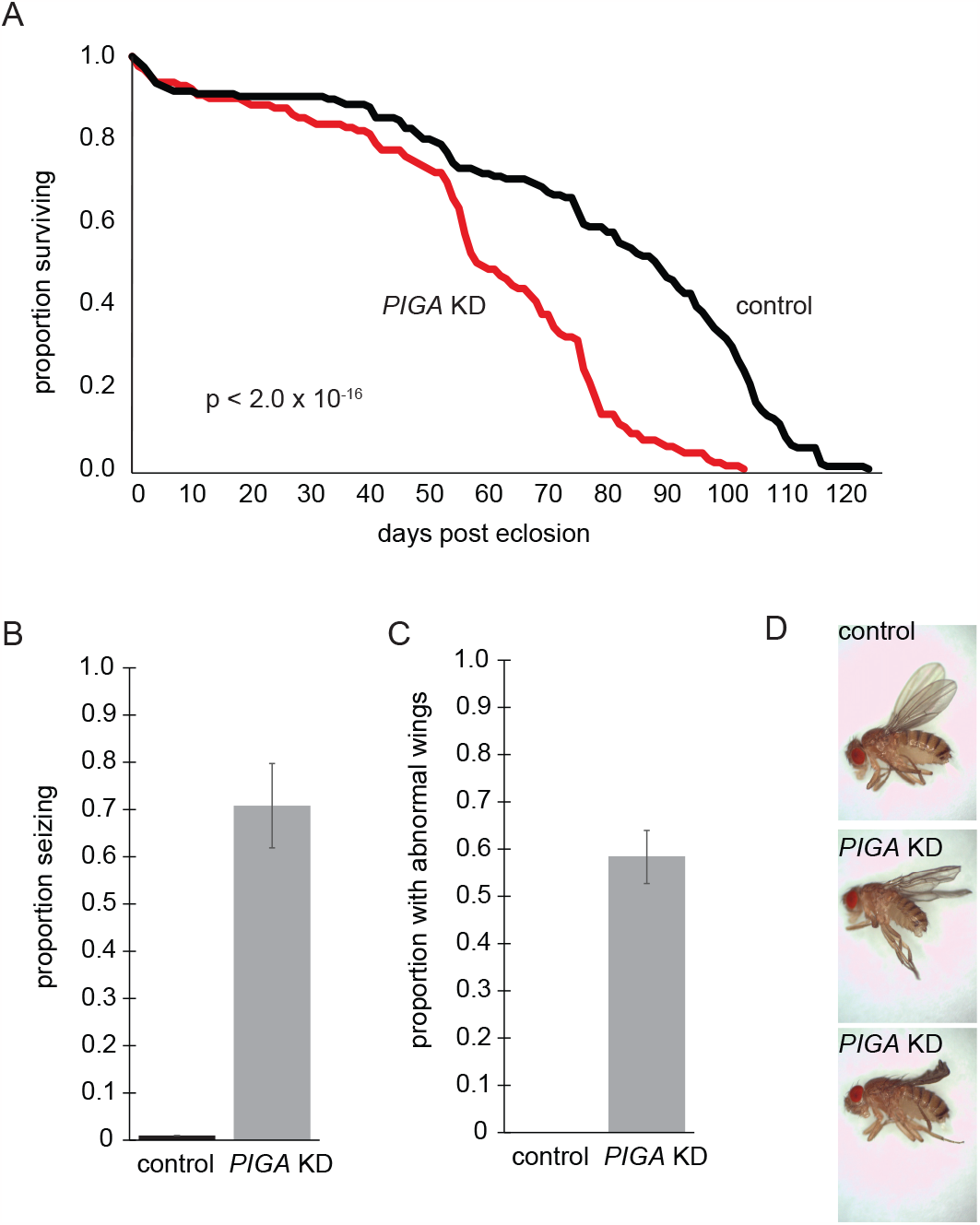
Phenotypes associated with glia-specific knockdown of *PIG-A*. A) Flies with glia-specific knockdown of *PIG-A* have reduced lifespan compared to control wildtype flies. B) Nearly 70% of glia-specific knockdown flies have seizures, and C) nearly 60% have abnormal, crumpled wings. D) Examples of crumpled wings in glia-specific knockdown flies compared to control wildtype flies. control = wildtype; *PIG-A* KD = glia-specific knockdown

### Transcriptomic data

To determine how loss of *PIG-A* in neurons vs. glia affects gene expression in the brain, we performed RNAseq analyses on whole heads. Neuron-specific knockdown heads showed a modest change in expression with 191 genes upregulated and 104 genes downregulated compared to control heads (Figure 4A; Table S1). Glia-specific knockdown heads showed a more substantial change in expression with 493 genes upregulated and 257 genes downregulated compared to control heads (Figure 4B; Table S1).

**Figure 4.**
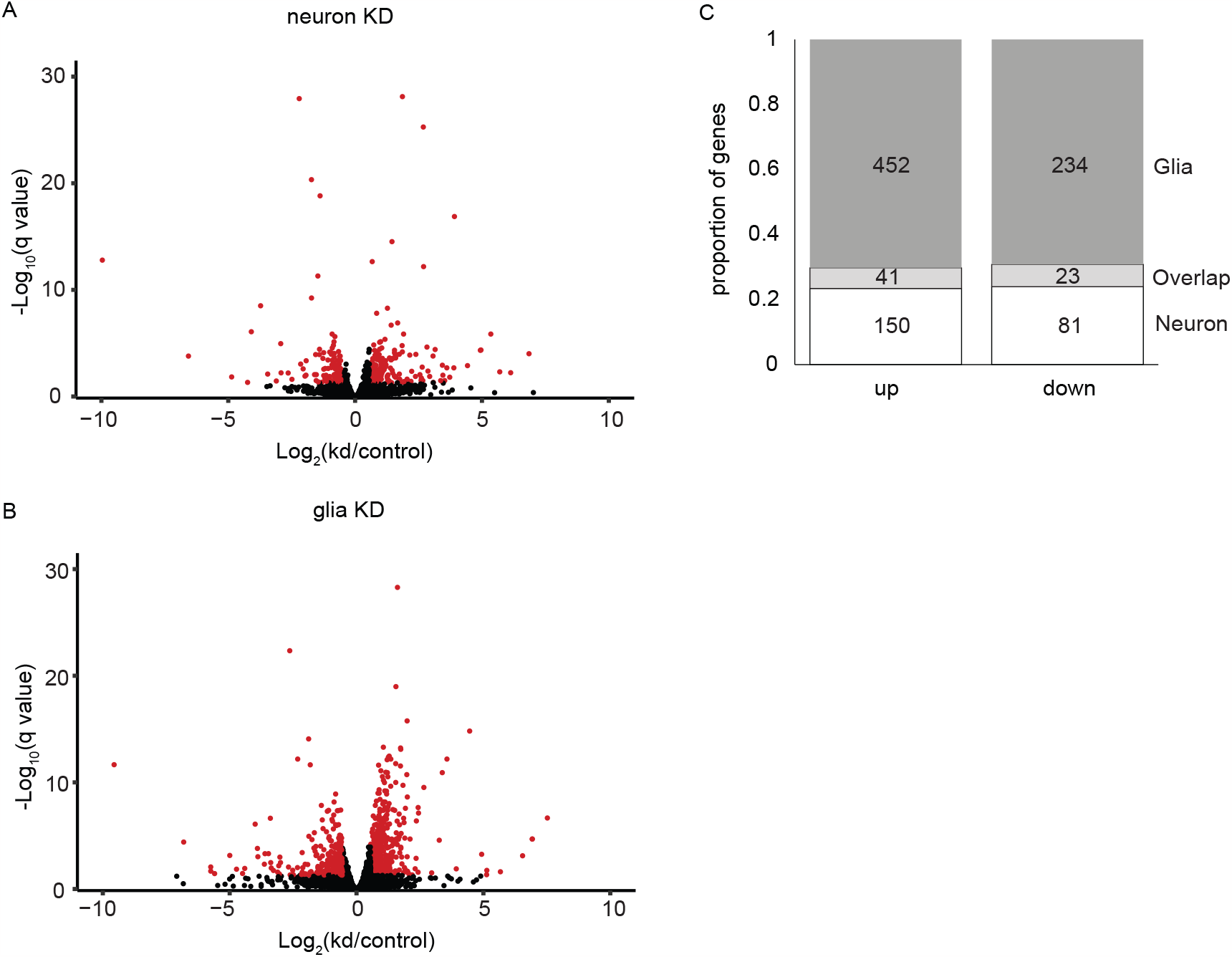
RNAseq results for neuron- and glia-specific knockdown of *PIG-A*. Volcano plots of A) neuron- and B) glia-specific knockdown of *PIG-A*. Red indicates transcripts that are ≥1.5 fold up or downregulated with q value ≤ 0.05. C) Proportion of genes that are uniquely or commonly up and down regulated in neuron- and glia-specific knockdown of *PIG-A*.

Gene ontology (GO) analyses of genes upregulated (≥1.5 fold; padj < 0.05) in the neuron-specific knockdown (Figure 5A; Table S2) showed a very strong enrichment for categories involved in glucose metabolism and glycolysis. For example, some of the most enriched categories include monosaccharide biosynthetic process (GO:0046364; fold enrichment: 35.3; p < 3.08 × 10^−4^), glucose 6-phosphate metabolic process (GO:0051156; fold enrichment: 29.9; p < 5.09 × 10^−4^), glucose metabolic process (GO:0006006; fold enrichment: 23.0; p < 5.70 × 10^−6^), glycolytic process (GO:0006096; fold enrichment: 18.6; p < 4.89 × 10^−4^), and hexose metabolic process (GO:0019318; fold enrichment: 18.6; p < 1.27 × 10^−7^). This suggests that there is a substantial increase in glycolysis in brains when PIGA function is eliminated in neurons. In fact, nearly all components of the glycolysis pathway are consistently 1.5-2.0-fold upregulated (FIGURE 6). We also observed enrichment for several categories involved in amino acid metabolism, including serine family amino acid metabolic process (GO:0009069; fold enrichment: 25.9; p < 7.74 × 10^−4^), alpha-amino acid catabolic process (GO:1901606; fold enrichment: 13.2; p < 3.14 × 10^−5^), and many others. Products of amino acid catabolism are often used in glycolysis. Despite this overwhelming increase in glycolysis-related gene expression, only two components of the TCA cycle are up regulated, suggesting that this increase in glycolysis is not related to changes in energy metabolism.

**Figure 5.**
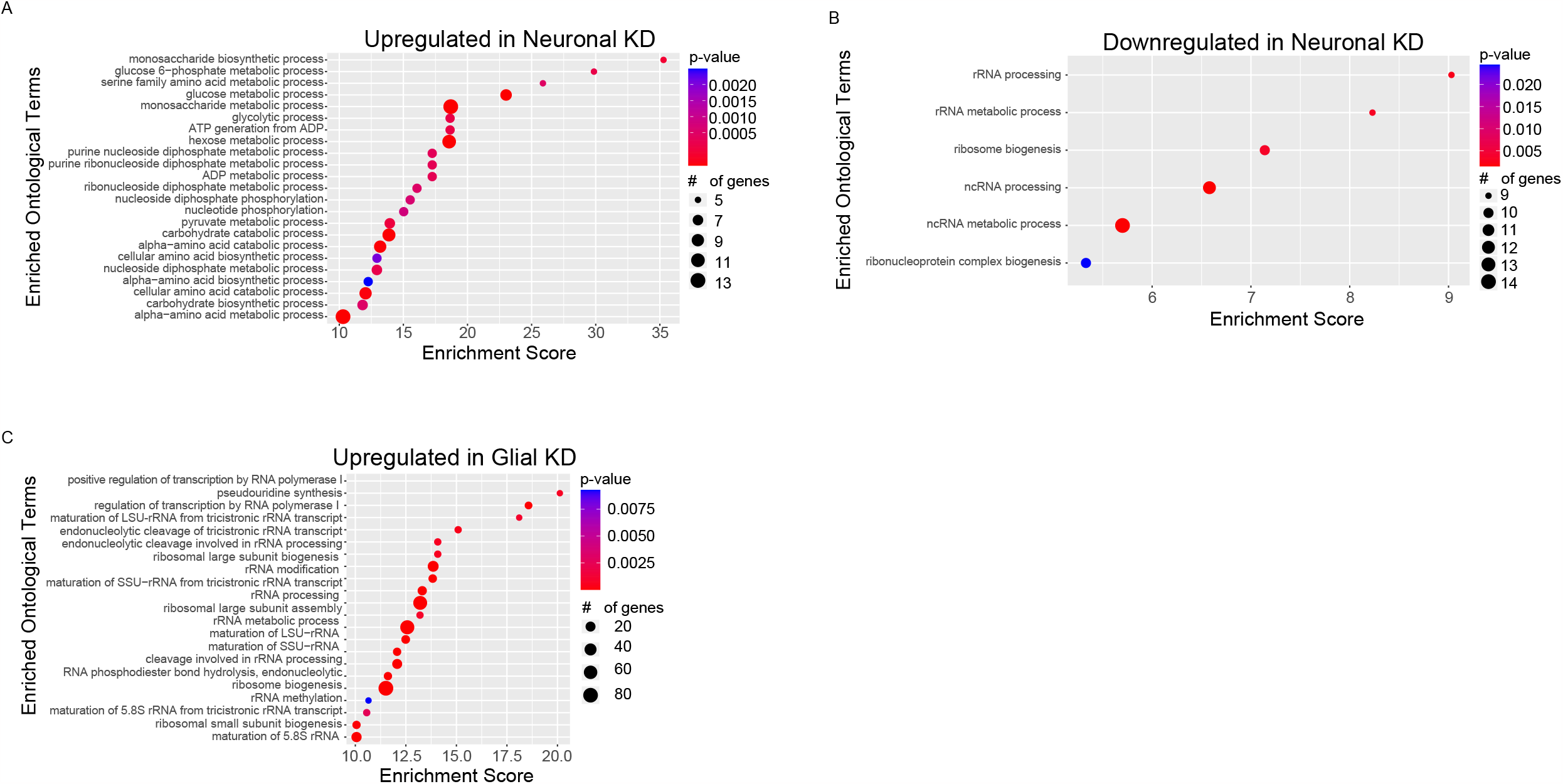
GO analyses of differentially expressed genes. Enriched GO terms for genes A) upregulated and B) downregulated in neuron-specific knockdown of PIG-A, and C) downregulated in glia-specific knockdown. There was no enrichment in genes upregulated in glia-specific knockdown.

**Figure 6.**
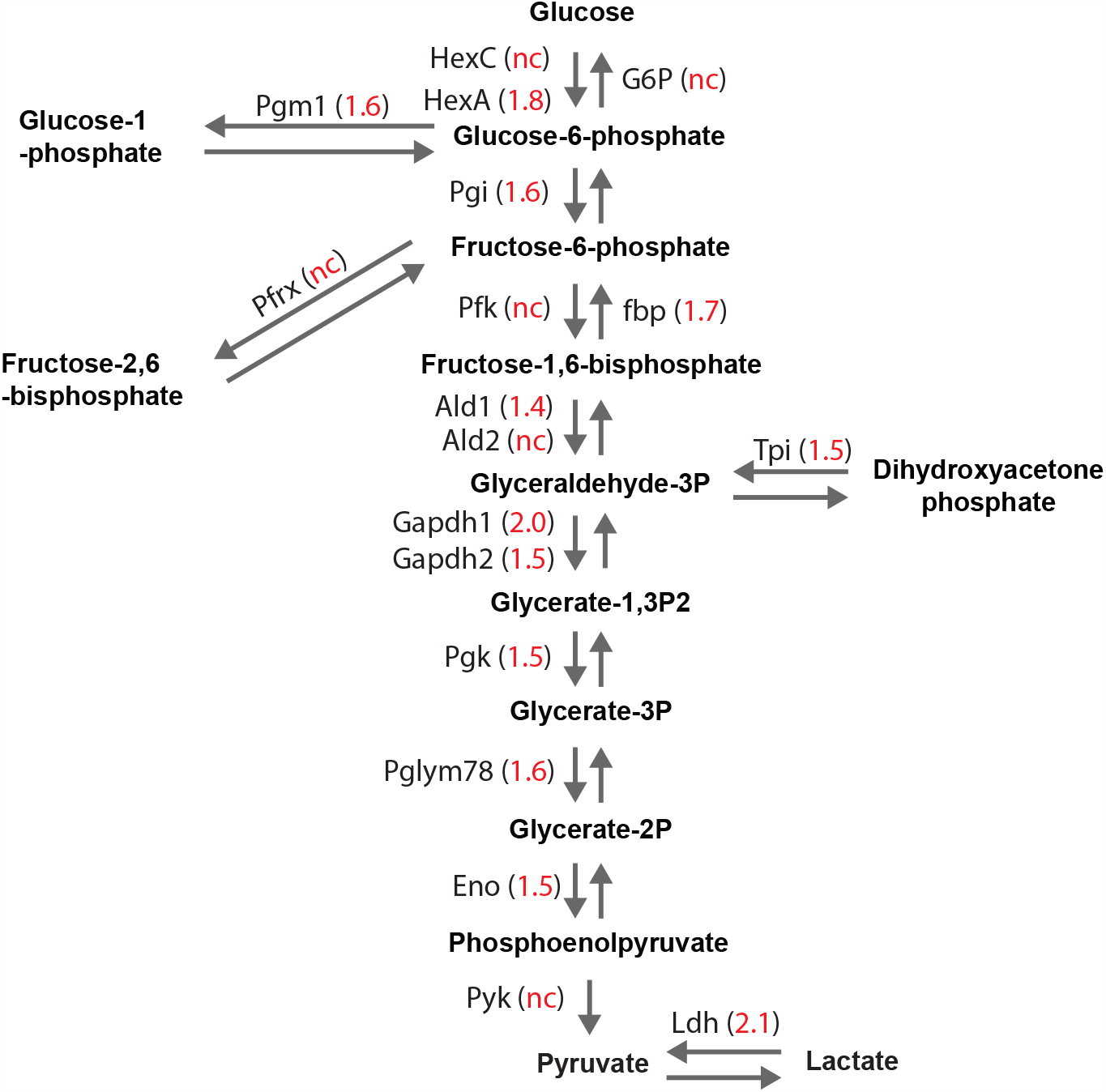
The glycolysis pathway is upregulated in neuron-specific knockdown of *PIG-A*. Nearly all the components of the glycolysis pathway are up regulated in neuron-specific knockdown of *PIG-A*. Numbers indicate fold change. nc = not changed.

GO analyses of genes downregulated (≤1.5 fold; padj < 0.05) in the neuron-specific knockdown (Figure 5B; Table S2) showed enrichment for genes involved in ribosome biogenesis: rRNA processing (GO:0006364; fold enrichment: 9.0; p < 2.58 × 10^−3^), ribosome biogenesis (GO:0042254; fold enrichment: 7.14; p < 3.61 × 10^−3^), ncRNA processing (GO:0034470; fold enrichment: 6.58; p < 1.45 × 10^−4^), ribonucleoprotein complex biogenesis (GO:0022613; fold enrichment: 5.33; p < 2.45 × 10^−2^), and other related functions. However, these genes encode mostly nucleolar proteins and most canonical ribosome biogenesis genes are unchanged. In fact, we see that these downregulated genes are enriched for the nucleolus cellular compartment (GO:0005730; fold enrichment: 11.63; p < 2.93 × 10^−3^). A survey of all downregulated genes did not identify any genes with functions related to glucose metabolism found in the upregulated gene set.

GO analyses of the genes upregulated (≥1.5 fold; padj < 0.05) in the glia-specific knockdown (Figure 5C; Table S2) also showed enrichment for ribosome biogenesis (GO:0042254; fold enrichment: 11.5; p < 2.32 × 10^−50^), rRNA processing (GO:0006364; fold enrichment: 13.2; p < 7.05 × 10^−44^), ncRNA processing (GO:0034470; fold enrichment: 8.3; p < 3.70 × 10^−39^), and ribosomal large subunit biogenesis (GO:0042273; fold enrichment: 13.9; p < 7.50 × 10^−18^). Unlike genes downregulated in the neuronal knockdown, this enrichment includes >80 genes directly involved in ribosome biogenesis. We also found enrichment in amino acid metabolism genes including pseudouridine synthesis (GO:0001522; fold enrichment: 18.6; p < 2.97 × 10^−5^), arginine metabolic process (GO:0006525; fold enrichment: 17.2; p < 3.51 × 10^−2^), and S-adenosylmethionine metabolic process (GO:0046500; fold enrichment: 17.2; p < 3.45 × 10^−2^). We also observed enrichment in components of the SRP-dependent cotranslational protein targeting to membrane, translocation (GO:0006616; fold enrichment: 17.2; p < 3.56 × 10^−2^), including nearly all the components of the Sec61 translocon and signal recognition particle receptor. Upon manual inspection of these genes, we found >30 genes that encode ER resident proteins including BiP and HYOU1, both involved in folding and processing nascent polypeptides that are translated into the ER through the translocon. Together, these data suggest that loss of PIGA in glia results in a large upregulation of genes in the brain involved in producing proteins that are processed through the ER, like GPI anchor proteins. GO analyses of genes downregulated (≤1.5 fold; padj < 0.05) in the glia-specific knockdown showed no functional enrichment.

We next examined the overlapping transcriptomic profile between neurons and glia (Figure 4C; Table S3). Forty-one genes are commonly upregulated at least 1.5-fold, between the two datasets. GO analyses of these common upregulated genes identified alpha-amino acid catabolic process (GO:1901606; fold enrichment: 36.2; p < 3.58 × 10^−4^) and cellular amino acid catabolic process (GO:0009063; fold enrichment: 33.1; p < 4.84 × 10^−2^). Each independent set of upregulated genes also showed upregulation for amino acid metabolism related functions. There is strong correlation in the degree of upregulation between these overlapping genes (r^2^ = 0.90; p = 1.3x10^−15^). We expanded the analyses to genes that were up regulated in either data set with a more relaxed 1.25-fold up regulation in the other data set to look more broadly at overlapping upregulated genes. While amino acid metabolism-related functions remain, we find enrichment for SRP-dependent cotranslational protein targeting to membrane, translocation (GO:0006616; fold enrichment: 35.5; p < 5.31 × 10^−2^) and hexose metabolic process (GO:0019318; fold enrichment: 10.1; p < 2.0 × 10^−2^). Each of these was identified in glia and neurons, respectively, but not the other. The degree of upregulation between these genes is still positive between neurons and glia (r^2^ = 0.56; p = 5.3x10^−16^).

There are only 23 genes that are commonly downregulated at least 1.5-fold (Figure 4C; Table S3). These commonly downregulated genes show strong correlation (r^2^ = 0.96; p = 2.6x10^−13^). There are 77 genes that are commonly downregulated at the more relaxed cutoff of 1.25-fold change. These genes are also strongly correlated (r^2^ = 0.71; p = 5.1x10^−13^). There is no functional enrichment in either group.

## DISCUSSION

Here we report the first *Drosophila* model of a disorder of GPI anchor biosynthesis. PIGA is the first step in GPI anchor synthesis and causes neurodevelopmental delay, movement disorder, and epilepsy (BAYAT *et al*. 2020). Using various genetic techniques, we show that we can model each of these phenotypes in *Drosophila*. In many ways, the *Drosophila* models mirror what has been reported for mouse models of PIGA-CDG, but we also are able to model certain patient phenotypes that have not been reported in mice.

As expected, complete loss of PIGA function is larval lethal in *Drosophila*. It is unlikely that proper GPI anchor biosynthesis can occur without PIGA function. In line with this, complete PIGa knockout in mice are early embryonic lethal. Further, no PIGA-CDG patients have been reported with predicted null mutations. While a few protein truncating mutations have been reported, they all produce truncated mutant protein. It was surprising that the *Drosophila* null mutants survived through 3^rd^ instar larval development. We suspect that this was likely possible because of maternal deposition of PIGA protein itself and a variety of GPI anchored proteins required for proper development. More work is needed to determine if this is the case.

Heterozygous null *Drosophila* with 50% function have similar amount of activity that is predicted in PIGA-CDG patients. Because *PIGA* is X-linked in humans, only males are affected, and they carry partial loss-of-function alleles that are predicted to have <50% function. Strikingly, these heterozygous null *PIG-A Drosophila* show a strong seizure phenotype that is very easily elicited. We noticed in our analyses that these flies would show seizure behavior if their vials were moved or tapped. Upon proper seizure analysis with the bang sensitive test, we found that about a third of the flies show strong seizure behavior. We think that this is an underestimation, as it is likely that many of the flies are in a refractory period. While we were not able to test if movement or other neurological phenotypes were normal, we did note that when flies are not seizing, they are climbing normally like wildtype flies. The ease at which these heterozygous flies have seizures matches a recent description of mosaic *PIGA* heterozygous knockout female mice (X linked, but one copy is randomly inactivated) that appear to also have easily elicited seizures when they fall (LUKACS *et al*. 2020). These heterozygous null flies will be a useful tool for studying the pathogenesis of seizures in PIGA-CDG and for testing possible small molecule therapeutics.

Because of the lethality and the inability to probe for other phenotypes in the homozygous null flies, we opted to use RNAi against PIG-A for further analyses. While knockdown by this RNAi was >60%, it is difficult to compare this to the heterozygous null, as qPCR was performed on whole larvae, a mixture of different cells that likely have different knockdown efficiencies. However, the fact that ubiquitous knockdown resulted in similar lethality as the null, suggests that at least for the critical cells, RNAi is quite effective. To determine if we could model other PIGA-CDG phenotypes in the fly, we generated neuron- and glia-specific knockdowns. Neuron-specific knockdown generated flies with both a movement disorder and neuromuscular defects, like what is observed in PIGA-CDG (BAYAT *et al*. 2020). Strikingly, neuron-specific knockdown did not produce observable seizures. We tried a number of different ways to elicit seizures beyond the bang sensitivity test and were not able to observe any seizure activity. Glial-specific knockdown generated flies with a profound seizure phenotype, much more severe than the heterozygous null flies. Glial-specific knockdown resulted in flies that likely have spontaneous seizures. The majority of flies seize upon bang sensitivity testing. However, we noticed that many of the flies will drop and seize even if a lab member walks by the vial. This separation of patient-relevant phenotypes in different cell types suggests that PIGA -CDG has a complex etiology and that different cell types have different requirements for GPI anchor biosynthesis.

To begin to understand some of the molecular consequences of losing PIGA function, we performed RNAseq on heads from neuron- and glia-specific knockdown flies. While this was an imperfect experiment because the transcriptome signatures are from a mix of normal and knockdown cells, we still observed very strong, specific changes that are informative. It should be noted that because we used RNAi, the transcriptome responses reflect partial loss of PIGA function, rather than complete loss of function. Residual function more closely reflects what is observed in PIGA-CDG. Overall, glia-specific knockdown generated a larger change in the transcriptome, as compared to the neuron-specific knockdown. Perhaps this makes sense, as the glial-specific knockdown flies are phenotypically much sicker than the neuron-specific knockdown flies. The transcriptomes from these two models produced very specific, but unique patterns. When we knockdown PIG-A in neurons, we find a near uniform upregulation of enzymes in the entire glycolysis pathway. We also found upregulation of enzymes responsible for amino acid metabolism. It is likely that this is not an increase in energy production, as enzymes in the TCA cycle remain mostly unchanged. Rather, we think that the cells are responding to a loss of GPI anchor production. Many products of glycolysis and amino acid metabolism are the substrates for the PIGA-catalyzed synthesis of the UDP-GlcNAc precursor and GlcNAc-PI (CHIARADONNA *et al*. 2018). In fact, levels of UDP-GlcNAc act as a “sensor” to regulate glycolysis and other metabolic pathways (CHIARADONNA *et al*. 2018). This suggests that when neurons have reduced PIGA function, cells in the brain are upregulating pathways that may provide the precursor molecules needed to ramp up GPI anchor biosynthesis.

The larger transcriptional changes observed in the glial-specific knockdown flies suggest that cells are ramping up the production of protein translational and folding machinery. It appears that the cells are responding to a reduction in GPI anchored protein production and increasing machinery to produce more GPI anchored proteins. Nearly all the components of ER associated translation are upregulated, including the SRP complex and the entire SEC21 translocon complex. Further, many genes encoding ER resident proteins involved in folding processing newly translated proteins are upregulated. This does not appear to be a response to ER stress, as other classic ER stress response genes, including those involved in ERAD, are unchanged. When glia have reduced PIGA function, cells in the brain respond by increasing ER-associated translation.

It is striking that the transcriptomes of neuron- and glia-specific knockdowns are so different. To determine if there were any commonalties, we compared the transcriptome changes. There were very few genes that showed the same direction of change between the two data sets. But the genes in common were very strongly corelated, suggesting a core response. The top ten genes in this overlapping set all show at least four-fold change in expression. However, these strong response genes mostly have unknown function with no human orthologs. We observed that the commonly upregulated genes show enrichment for amino acid metabolism. This enrichment is also observed in the neuron-specific knockdown, but not the glia. Upon manual curation, many of the genes that are commonly upregulated are enzymes related to energy metabolism, which again, makes sense for the neuron-specific knockdown. However, it suggests that there may also be changes related to metabolism and glycolysis in the glia specific knockdowns as well, but does not reach the enrichment observed in the neuron-specific knockdown. It needs to be noted that all the major glycolysis enzymes are unchanged in the glia-specific knockdown. More work is needed to understand the cellular origin of the changes uncovered by RNAseq. Future analyses involving metabolomics and proteomics will help to elucidate the changes that occur when the GPI anchor synthesis is disrupted.

In this study, we report the initial analyses of the first PIGA-CDG models in *Drosophila*. This study reinforces that *Drosophila* can serve as an efficient tool to accurately model CDGs and other rare neurodevelopmental disorders. The data here set the stage for future modifier and drug screen studies to help identify potential therapeutic strategies for PIGA-CDG.

---

**Figure S1:** qRT-PCR for PIG-A transcript levels in the RNAi knockdown.

**Table S1:** RNAseq data for neuron- and glia-specific knockdown of PIG-A.

**Table S2:** GO analyses of RNAseq results.

**Table S3:** Genes that are up or downregulated in neuron, glia, or both.

## Acknowledgements

We thank Drs. Hugo Bellen and Oguz Kanka for generating and sharing the *PIG-A* null allele. We thank members of the Chow lab for helpful comments on this manuscript.

## Funding

This research was supported by an NIH NIGMS R35 award (R35GM124780) and a grant from the Primary Children’s Hospital Center for Personalized Medicine to CYC. KGO and MCA were supported through the NIH NIGMS T32 Fellowship from the University of Utah (T32GM007464). Stocks obtained from the Bloomington Drosophila Stock Center (NIH P40OD018537) were used in this study. MH was supported by a predoctoral fellowship from the American Epilepsy Society. HJT was supported by the NIH NIDDK T32 Fellowship from the University of Utah (T32DK110966).

## Notes

### Competing Interest Statement

The authors have declared no competing interest.

