## Supplementary figures and images for "*Drosophila* models of PIGA-CDG mirror patient phenotypes"

### Figure S1

Supplemental figure 1

Figure S1

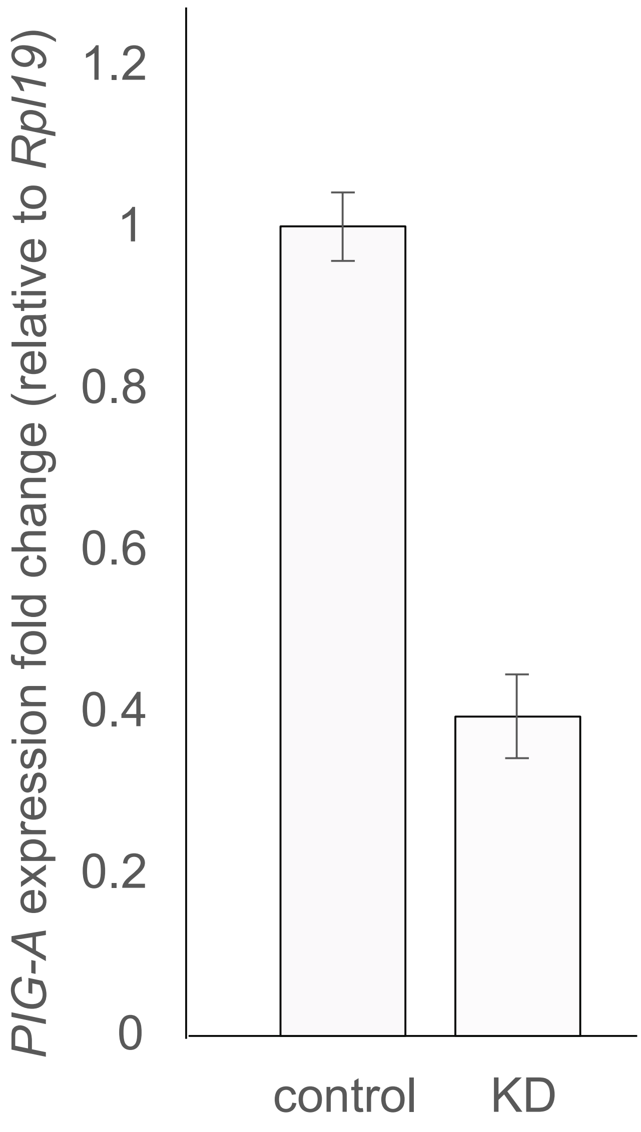
